# Differential Regulation of Rice Transcriptome to *Rhizoctonia solani* infection

**DOI:** 10.1101/2021.05.05.442799

**Authors:** Akash Das, Moin Mazahar, Ankur Sahu, Mrinmoy Kshattry, P.B. Kirti, Pankaj Barah

**Affiliations:** Department of Molecular Biology and Biotechnology, Tezpur University, Assam-784028, India; Department of Biotechnology, Indian Institute of Rice research, Hyderabad- 500030, India; Leibniz Institute of Plant Genetics and Crop Plant Research (IPK), Gatersleben Seeland- 06466, Germany; Agri Biotech Foundation, PJTS Agricultural University Campus, Hyderabad- 500030, India

**Keywords:** Systems Biology, RNA-Seq, Machine Learning, Transcriptomics, Rice, *Rhizoctonia solani*, Sheath Blight, Transcriptional Regulatory Networks

## Abstract

Sheath Blight (SB) disease in rice crop caused by the infection of the fungal pathogen *Rhizoctonia solani* (*R. solani*) is one of the severe rice diseases that can cause up to 50% yield losses. Naturally occurring rice varieties resistant to SB have not been reported yet. We have performed a Time-Series RNA-Seq analysis on a widely cultivated rice variety BPT-5204 for identifying its transcriptomic response signatures to *R. solani* infection at 1^st^, 2^nd^ and 5^th^ day post inoculation (dpi). In total, 428, 3225 and 1225 genes were differentially expressed in the treated rice plants post 1, 2 and 5 dpi, respectively. GO and KEGG enrichment analysis identified significant processes and pathways differentially altered in the rice plant after the fungal infection. Machine learning and network based integrative approach was used to construct Transcriptional Regulatory Networks (TRNs) of the rice plant at the three Time Points. Regulatory network analysis identified SUB1B, MYB30 and CCA1 as important regulatory hub Transcription Factors in rice during *R. solani* infection. Jasmonic acid signaling pathway was activated and in contrast, photosynthesis and carbon fixation processes were significantly compromised. Involvement of MAPK, CYPs, Peroxidases and PAL genes was observed in response to the fungal infection. Circadian clock was also strongly influenced by *R. solani* infection. Our integrative analysis identified 7 putative SB resistant genes altered in rice after *R. solani* infection and provided a better understanding of rice plant response to *R. solani* infection.

**One sentence summary:** Time series expression analysis of rice variety BPT-5204 identifies key molecular signatures involved in rice plant response to *R. solani* infection.

## INTRODUCTION

Rice Sheath Blight (SB) disease is one among the top three most devastating diseases of rice worldwide. It can cause up to 50 % of yield losses under optimal environmental conditions (Qingzhong et al., 2001). This disease is caused by a wide host range fungal pathogen, *Rhizoctonia solani*. The fungus is a soil borne necrotroph, divided into 14 anastomical groups (AG1 to AG13) based on the diversity in colony morphology, biochemical and molecular markers, pathogenicity, and aggressiveness. Subgroup AG1-1A is responsible for causing Sheath blight disease (Yuan et al., 2018). The primary source of inoculum is the load of sclerotia left over from the previous harvest, which later affects the forthcoming season upon activation by optimum environmental conditions. Contaminated water transmission for irrigation purposes paves the way for greater spread from one field to the others (Kobayashi et al., 1997). Unfortunately, naturally occurring rice germplasm showing absolute resistance to this catastrophic disease does not exist and the control of the disease lies solely in the use of appropriate fungicides.

The rapid increase in population and global climate change affect the food security demands, which necessitate immediate attention. Predictions are made that rice prices can surge by 37% by the year 2050 as a result of climate change that results in significant reduction in total rice yields by 14, 10 and 15% in South Asia, East Asia and Sub-Saharan Africa, respectively (Research Institute (IFPRI), 2009). Plants being sessile organisms cannot migrate to circumvent the stress conditions and need to adapt themselves with adversities like rising temperature (Barah et al., 2013). Variations in temperature have direct correlation with plant disease by altering host resistance mechanisms of plants towards a particular disease (Peng et al., 2004). Evidence also suggests that SB disease severity in crops rapidly increases with elevated carbon dioxide levels (Luck et al., 2011). Hence, it forms a matter of serious concern to devise strategies to improve the crop varieties to sustain such environmental adversities.

In recent years, some studies have tried to delineate the mechanism of rice responses to the impending pathogen attack at a molecular level. Various signal transduction pathways have been shown to be involved in rice in response to infection by *R. solani*. These pathways include Jasmonic acid, Octadecanoid, Ethylene and Salicylic Acid (SA) and Calcium/ Calmodulin pathways. Studies show that Jasmonic acid (JA), Lipoxygenase (LOX) and Octadecanoid pathways are induced upon *R. solani* infection (Taheri et al., 2010). Mutant rice deficient in JA responses showed enhanced susceptibility to SB pathogen indicating that JA pathway plays a crucial role in rice resistance to *R. solani*. LOX is a precursor for the JA pathway and the corresponding gene is shown to be overexpressed in rice under *R. solani* infected conditions (Sayari et al., 2014). Substantial evidence also indicated the involvement of ethylene (ET) pathway in resistance of rice against SB. Rice overexpressing ET biosynthetic genes tend to increase the expression of PR1b and PR5 genes, which results in resistance (Helliwell et al., 2013). The induction of PR genes is an indication of *Systemic Acquired Resistance (SAR)* activation in plants. Further, active involvement of SAR pathway is also suggested by the enhanced expression of PR-5, PR-3, PR-9, PR12, PR-13 and Phenylalanine Ammonia Lyase (PAL) genes. PAL in an essential SAR marker (Sayari et al., 2014). Calcium and calmodulin signaling pathways play crucial roles in rice stress responses. Various genes including OsCam1– 1 belonging to calcium calmodulin signaling pathway showed enhanced expression in *R. solani* infected rice suggesting its crucial role in defense (Zhang et al., 2017).

Systems biology is often defined as a global approach for developing comprehensive understanding of Plant-Pathogen interactions by establishing relationships between data in the form of genes, RNA, proteins and metabolites (Mishra et al., 2019). Rapid development and gradual reduction in cost of next generation sequencing (NGS) technologies paved the way for methods like RNA-Sequencing to conduct genome scale experiments for studying plant pathogen interactions. Very few studies using RNA-Seq based approaches have been used to determine molecular signatures related to *R. solani* infection in different susceptible and resistant rice varieties (Ghosh et al., 2017; Zhang et al., 2017; Shi et al., 2020). Using microarray analysis, Venu *et*.*al* have identified both up-regulated and down-regulated rice genes after SB disease condition (Venu et al., 2007). Comparative RNA-Seq analysis predicted rice transcriptome profiles for Lemont (susceptible) and TeQing (Moderately resistant) varieties in SB disease condition (Zhang et al., 2017). The experiment identified a total of 4806 differentially expressed genes (DEGs) and suggested photorespiration, photosynthesis and JA pathway to have major roles in rice resistance mechanism to the disease. Another similar analysis was conducted on two different varieties of rice-Yanhui-888 (YH), a moderately resistant cultivar and Jingang-30 (JG), a susceptible cultivar to detect rice responses against *R. solani* infection (Shi et al., 2020). This study identified 3085 and 285 DEGs in JG and YH rice varieties, and showed that EIN2, WRKY33 genes are important in rice responses to *R. solani*.

Karmakar *et.al*. (2019) conducted metabolome-transcriptome studies to identify metabolome and transcript level alteration in rice infected with *R. solani*. This study identified 38 differentially expressed proteins (DEPs) and 40 metabolites in rice after *R. solani* infection. Study also suggests carbohydrate metabolism to be a crucial point of focus for rice under infected conditions (Karmakar et al., 2019). However, the exact molecular crosstalk mechanism still remains unclear and the search for naturally occurring resistant germplasm continues.

To this end, a comparative Time-Series based RNA-Seq analysis of a widely cultivated rice (BPT-5204) variety during *R. solani* infection was performed. Differentially regulated rice transcriptome was decoded in 1^st^, 2^nd^ and 5^th^ day post *R. solani* infection. Our study identifies key molecular signatures in terms of genes, pathways, biological processes and regulatory hub genes, which were significantly altered in rice plants after *R. solani* infection and might play a crucial role in plant defense mechanism. Additionally, Transcriptional Regulatory Networks provided insights into rewiring patterns of Transcription Factors (TFs) in all the three days after infection.

## MATERIALS AND METHODS

### Rice plant growth and inoculation

*Oryza sativa sp. indica* cultivar BPT-5204 (Samba Mahsuri) was used in this study. The sclerotia of *R. solani* were cultured on Potato Dextrose Agar (39 g/L) medium at 28 °C. Freshly grown equal-sized sclerotia blocks were placed on the leaf sheaths at the late tillering stage of the rice plants grown in warm and moist conditions (Prathi et al., 2018). After inoculation, the infected plants were maintained at 28 °C in dark for one to three days. After 1^st^, 2^nd^ and 5^th^ day of treatment, leaf tissue samples were collected as three biological replicates.

### RNA extraction and sequencing

RNA was extracted from 50-100 mg leaf tissue using Qiagen RNAeasy Plant Mini kit, according to the manufacturer’s instructions. Briefly, leaves were homogenized using liquid nitrogen and RLT buffer in TOMY microsmash homogenizer. The lysate was centrifuged to remove debris. Supernatant was mixed with equal volume of 70% ethanol and loaded onto Qiagen RNeasy column and further steps were followed as per manufacturer’s guidelines, including on-column DNase treatment using Qiagen RNase free DNase. RNA was eluted in Nuclease free water. The quantification, quality and integrity of the RNA was assessed using Nanodrop2000 (Thermo Scientific, USA), Qubit (Thermo Scientific, USA) and Bioanalyzer 2100 (Agilent, USA), respectively.

RNA sequencing libraries were prepared with Illumina-compatible NEBNext® Ultra™ II Directional RNA Library Prep Kit (New England BioLabs, MA, USA) at Genotypic Technology Pvt. Ltd., Bangalore, India. An aliquot of 500 ng of total RNA was taken for mRNA isolation, fragmentation and priming. Fragmented and primed mRNA was further subjected to first strand synthesis followed by second strand synthesis. The double stranded cDNA was purified using JetSeq Beads (Bioline, USA). Purified cDNA was end-repaired, adenylated and ligated to Illumina multiplex barcode adapters as per NEBNext® Ultra™ II Directional RNA Library Prep protocol followed by second strand excision using USER enzyme at 37 °C for 15mins. Illumina Universal Adapters were used in the study. Adapter ligated cDNA was purified using Jet Seq Beads and was subjected to 11 cycles of Indexing-PCR (98 °C for 30 sec, cycling (98 °C for 10sec, 65 °C for 75 sec) and 65 °C for 5 min) to enrich the adapter-ligated fragments. Final PCR product (sequencing library) was purified with Jet Seq Beads, followed by library quality control check. Illumina-compatible sequencing library was quantified by Qubit fluorometer (Thermo Fisher Scientific, MA, USA) and its fragment size distribution was analysed on Agilent 2100 Bioanalyzer. The libraries were pooled in equimolar quantities and sequenced on Illumina HiSeq X10 sequencer (Illumina, San Diego, USA) using 150 bp paired-end chemistry and HiSeq X10 SBS reagents. An average of 13.9±3.44 million reads were obtained for the samples.

### Quality check and trimming of reads

All reads were analyzed using robust and benchmarked in-house RNA-Seq pipeline (https://github.com/evolomics-group/rna-seq-pipeline). The pipeline being modular can be used in both plant and human model systems (Sahu et al., 2020; Sahu et al., 2020; Roy et al., 2020). Firstly, quality check for the raw fastq files was conducted using fastQC tool. Following criteria were considered to filter out the low-quality reads: reads lower than Q30 phred score; reads shorter than 15 bp; Illumina adapter clipping were discarded. These clean reads were then re-assessed for quality prior to alignment.

### Mapping and quantification of reads against the rice genome

Clean reads were mapped to rice reference genome Os-Nipponbare-Reference-IRGSP-1.0 from RAP-DB database (Sakai et al., 2013). Prior to mapping, indexing of the reference genome was done using the HISAT2 indexing scheme, which uses the Burrows-Wheeler transform and the Ferragina-Manzini (FM) as indexing parameters (Kim et al., 2015). Further, these clean reads were mapped against the reference index file using HISAT2. Samtools was used to convert the mapped output files (sam files) into binary files (bam files) (Li et al., 2009). The mapped binary files were quantified using the Subread package called featureCounts and counted at the feature (gene) level for further analysis (Liao et al., 2014).

### Normalization and visualization of count data

The R/Bioconductor package DESeq2 was implemented to perform the statistical analysis on count data (Love et al., 2014). Generalized linear model (GLM) and Wald test were used to evaluate the significance of differentially expressed genes (DEGs). Counts below a threshold of 10 were rejected. Benjamini-Hochberg multiple testing adjustment procedure was implemented to adjust the P Values. The rank score of DEGs was determined using lfcShrink function. Data transformation was done using variance stabilizing transformation. The amount of shrinkage was estimated using dispersion parameters for each gene. Principal component analysis was used to evaluate the overall effect of experimental covariates. Genes showing a significance of Padj value </= 0.05 were considered and termed as differentially expressed genes (DEGs).

### GO and pathway enrichment analysis

Gene Ontology (GO) annotations for all the day specific DEGs were performed using CARMO (Wang et al., 2015b). DEGs were classified according to their Biological Process (BP), Molecular Function (MF) and Cellular Component (CC) categories they are involved in. Only significant GO terms were ranked taking enrichment P Value (< 0.05) as cutoff. All data visualization and plotting were carried out in RStudio work environment. To check alterations occurring at the pathway level, the unique DEGs were also mapped against the Kyoto Encyclopedia of Genes and Genomes (KEGG) using open source and upgraded tool KOBAS 2.0 (Xie et al., 2011). Significant pathway terms were ranked taking enrichment P Value (< 0.05) as cutoff. Top 20 enriched KEGG pathways for all the three days were visualized as dot plots using ‘ggplot2’ package in RStudio.

### Transcription Factor identification and TF-TG network construction

Transcription Factors were extracted from the DEGs by intersecting the same with a list of 1862 TFs RAP id entries collected from PlantTFDB (Guo et al., 2007) database. TFs expressed as common in all the three days were identified and further visualized as heatmaps using the MeV tool. For construction of the Transcription Factor – Target Gene (TF-TG) networks, the upstream sequences of the day specific DEGs were initially extracted using RSAT (Thomas-Chollier et al., 2008). The region selected was between -1 kb and -1 of the TSS (transcription start site). PWMs for all the TFs were downloaded from the Cis-BP database (Weirauch et al., 2014). The downloaded PWMs were used to scan the upstream sequences of DEGs in FASTA format establishing relations between each TF and TG. The whole process was conducted using the FIMO of MEME suite under a High Performance Computing (HPC) environment with slurm acting as a queue manager for a streamlined workflow (Grant et al., 2011). Finally, TF-TG relations were visualized as networks using Cytoscape 3.8.1 (Su et al., 2014) for each day separately.

### PR genes study and benchmarking

A comprehensive and non-redundant list of 48 putative rice Resistant Genes (RGs) found in three major diseases of rice i.e., Bacterial Blight (BB), Rice Blast (RB) and Sheath Blight (SB) was prepared by integrating data collected from PRGdb 3.0 database (Osuna-Cruz et al., 2018) (for BB and RB disease) and manual curation of literature to find genes conferring resistance against SB (Supplemental Data Set 6). DEGs were compared with this list to check for the presence of RGs in our data and visualized using RStudio.

## RESULTS

### Transcriptome reprogramming of rice plants in response to *R. solani* infection

In the present study, we used a Time-Series RNA-Seq approach to decipher the differential regulation in the rice transcriptome profile upon *R. solani* infection. Rice cultivar BPT-5402 was inoculated with *R. solani* and samples were harvested at 1^st^, 2^nd^ and 5^th^ day post inoculation with the pathogen. A total of 428 (292 up-regulated, 136 down-regulated), 3225 (2102 up-regulated, 1123 down-regulated) and 1225 (739 up-regulated, 486 down-regulated) unique DEGs were identified in day 1, 2 and 5 samples, respectively in rice in response to *R. solani* (Fig. 1A) (Supplemental Data Set 1). Maximum number of rice genes were expressed in samples of day 2 infection followed by day 5 and day 1. Moreover, the number of up-regulated DEGs outnumbered the down-regulated DEGs at each of the Time Points. We further created two unified lists of the DEGs (combining all the 3 Time Points) into groups of up-regulated and down-regulated genes. A total of 2,456 and 1,515 DEGs were found to be only up-regulated and down-regulated respectively. In the list of 2,456 up-regulated DEGs, 133 were differentially expressed at all the three Time Points (Fig. 1B) whereas 20 of the 1,515 total down-regulated DEGs were differentially expressed at all the three Time Points (Fig. 1C).

**Figure 1.**
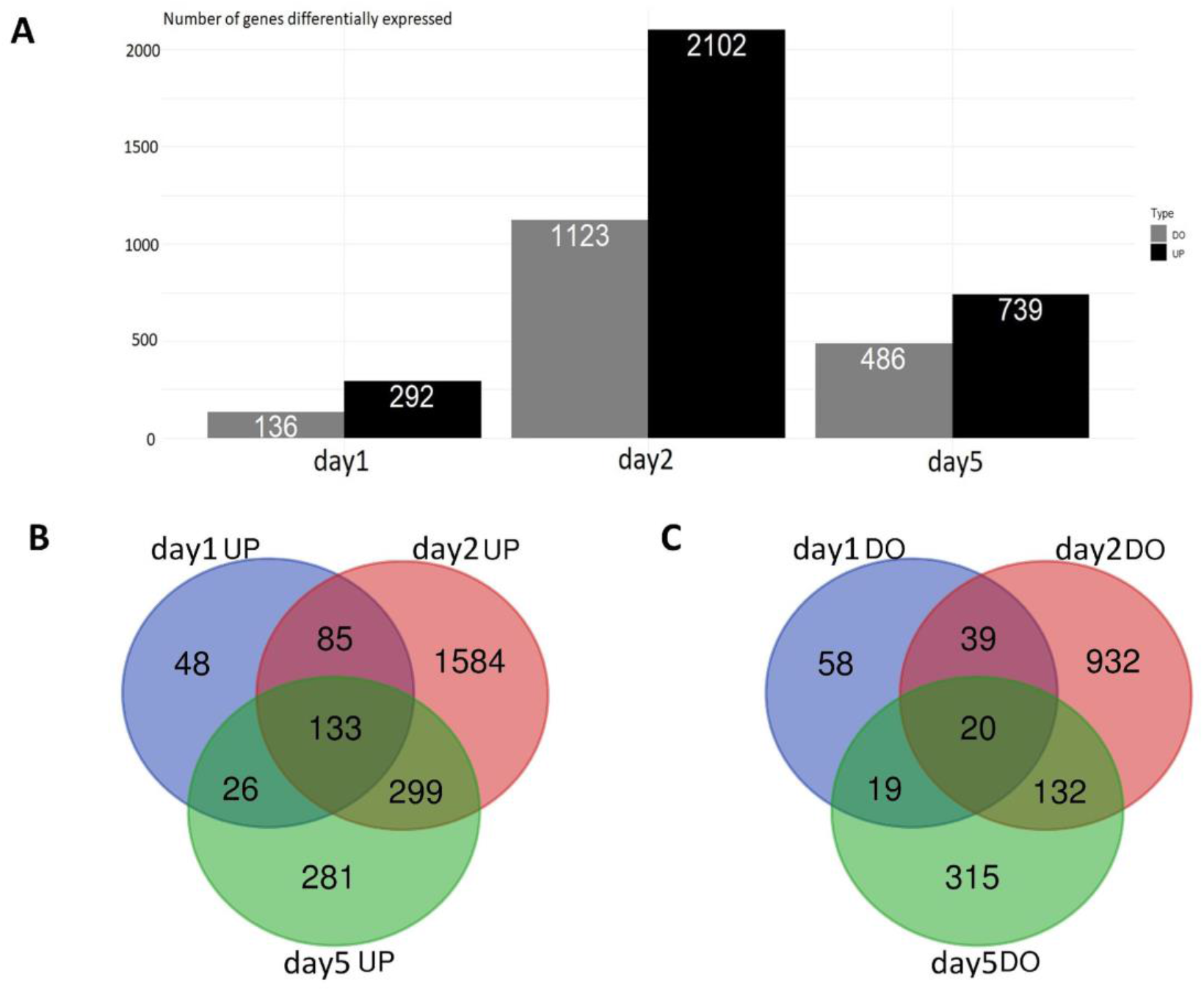
Differentially expressed genes (DEGs) in rice after *R. solani* infection. A, Bar plot showing the number of DEGs (up-regulated and down-regulated) identified in rice at 1^st^, 2^nd^ and 5^th^ day Time Points after *R. solani* infection. B, Venn diagram showing up-regulated DEGs expressed as common in day 1, 2 and 5 samples. A total of 133 genes were found to be up-regulated as common in all the three Time Points. C, Venn diagram showing down-regulated DEGs expressed as common in day 1, 2 and 5. A total of 20 genes were found to be down-regulated as common at all the three Time Points.

### Differential signatures of Biological Processes in Rice during *R. solani* infection

To elucidate the various biologically significant functions associated with the specific DEGs of day 1, 2 and 5 samples in rice after *R. solani* infection, we performed GO enrichment analysis using CARMO (Wang et al., 2015b). We had divided the DEGs into down-regulated and up-regulated lists for each of the Time Points and used them for GSEA (Gene Set Enrichment Analysis). Considering enrichment P Value threshold 0.05, we identified 38, 119 and 32 up-regulated and 23, 117 and 43 down-regulated GO terms belonging to Biological process (BP), Cellular Component (CC) and Molecular Function (MF) categories in day 1, 2 and 5 samples, respectively (Supplemental Data Set 2). Out of these enriched GO terms, we selected the top 20 enriched Biological Processes of day 1, 2 and 5 samples (up-regulated and/or down-regulated) for further analysis. The summary of the GSEA has been presented in Figure 2.

**Figure 2.**
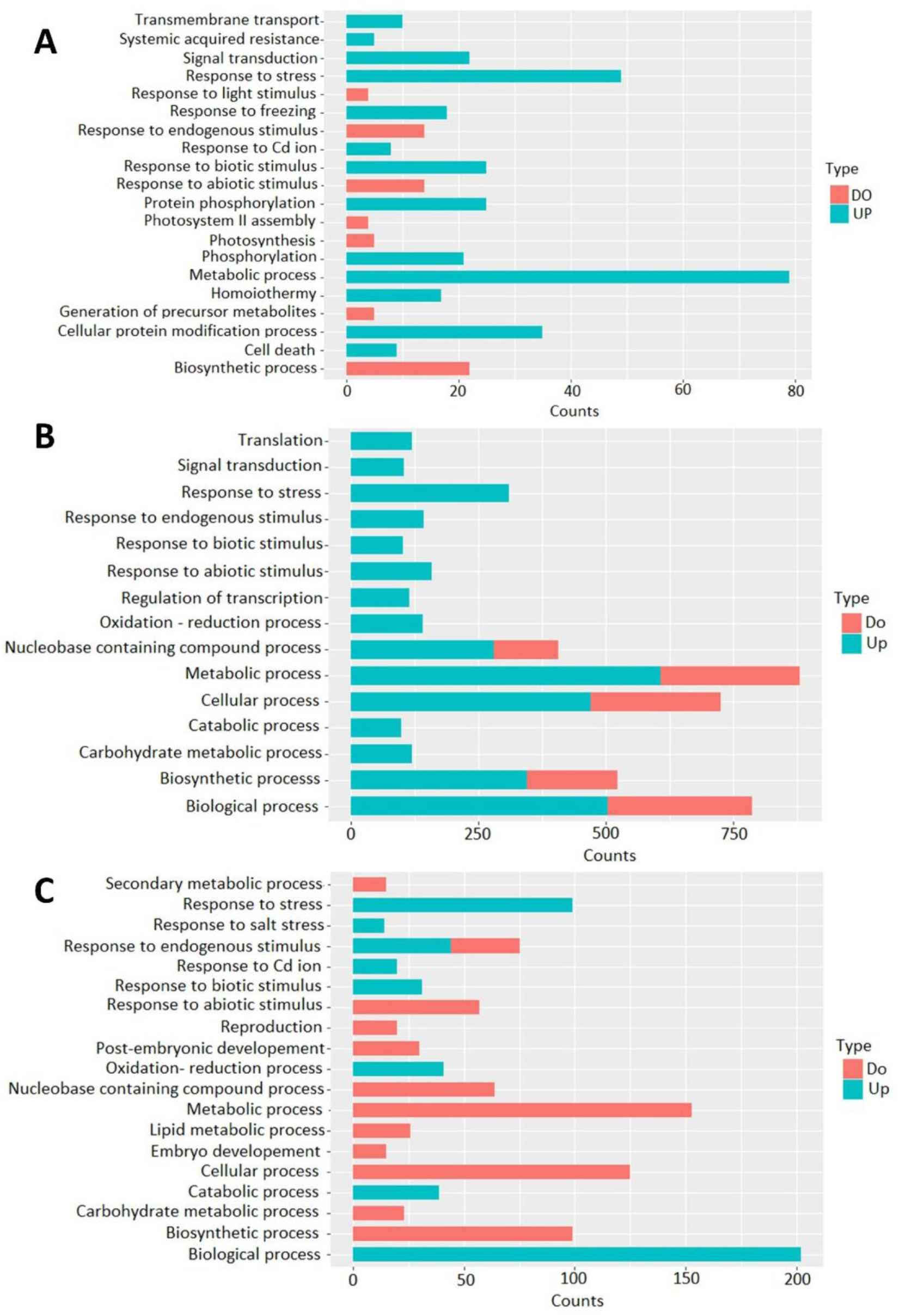
Gene Ontology (GO) analysis of significantly altered Biological Processes. Bar graphs showing the differential regulation of various Biological Processes and their expression type (UP= up-regulated and/or DO= down-regulated) in rice after *R. solani* infection at all the three Time Points i.e., day 1 (A), day2 (B)and day 5 (C).

To determine the highly significant Biological Processes associated with rice responses to *R. solani*, a comparison was carried out among the top 20 enriched BPs of all the three Time Points. The GO BP terms associated with response to biotic stimulus (GO:0009607), response to cadmium ion (GO:0046686), response to stress (GO:0006950), response to abiotic stimulus (GO:0009628) and biosynthetic process (GO:0009058) were found to be expressed as common among all the three Time Points. Interestingly, the BPs ‘response to biotic stimulus’, ‘response to cadmium ion’ and ‘response to stress’ were found to be only up-regulated at all the three Time Points. A step further, the genes belonging to each of the above-mentioned common BPs and differentially expressed at all the three Time Points have been listed in Supplemental Table 1. In summary, as the genes listed in Supplemental Table 1 belonging to common BPs were differentially expressed at all the three Time Points and were either up-regulated or down-regulated throughout the Time Points, they can be considered as potential *R. solani* responsive genes.

### Differential Pathway Enrichment analysis

In order to characterize the significant pathways that are differentially regulated in rice after *R. solani* infection, we performed pathway enrichment analysis from KEGG database using KOBAS (Xie et al., 2011). The KEGG pathway entries that were altered in rice after infection in day 1, 2 and 5 samples, respectively have been listed in Supplemental Data Set 3. Of them, the top 20 altered KEGG pathways (both up-regulated and down-regulated with enriched P Value threshold 0.05) enriched at all the three Time Points are represented as dot plots in Figure 3. An easy comparison of the top 20 significant KEGG pathways among the three Time Points was drawn (Supplemental Data Set 4). The pathway ‘Ribosome biogenesis in eukaryotes’ was found to be common and up-regulated in all the three samples, whereas ‘Metabolic pathways’ and ‘Biosynthesis of secondary metabolites’ appeared more as down-regulated in all the three samples. The gene ‘CYP51H4/ Os02g0323600’ (a Cytochrome P450-like protein) is a member of both the ‘Metabolic pathways’ and ‘Biosynthesis of secondary metabolites’ pathways and was found to be common for all the three samples.

**Figure 3.**
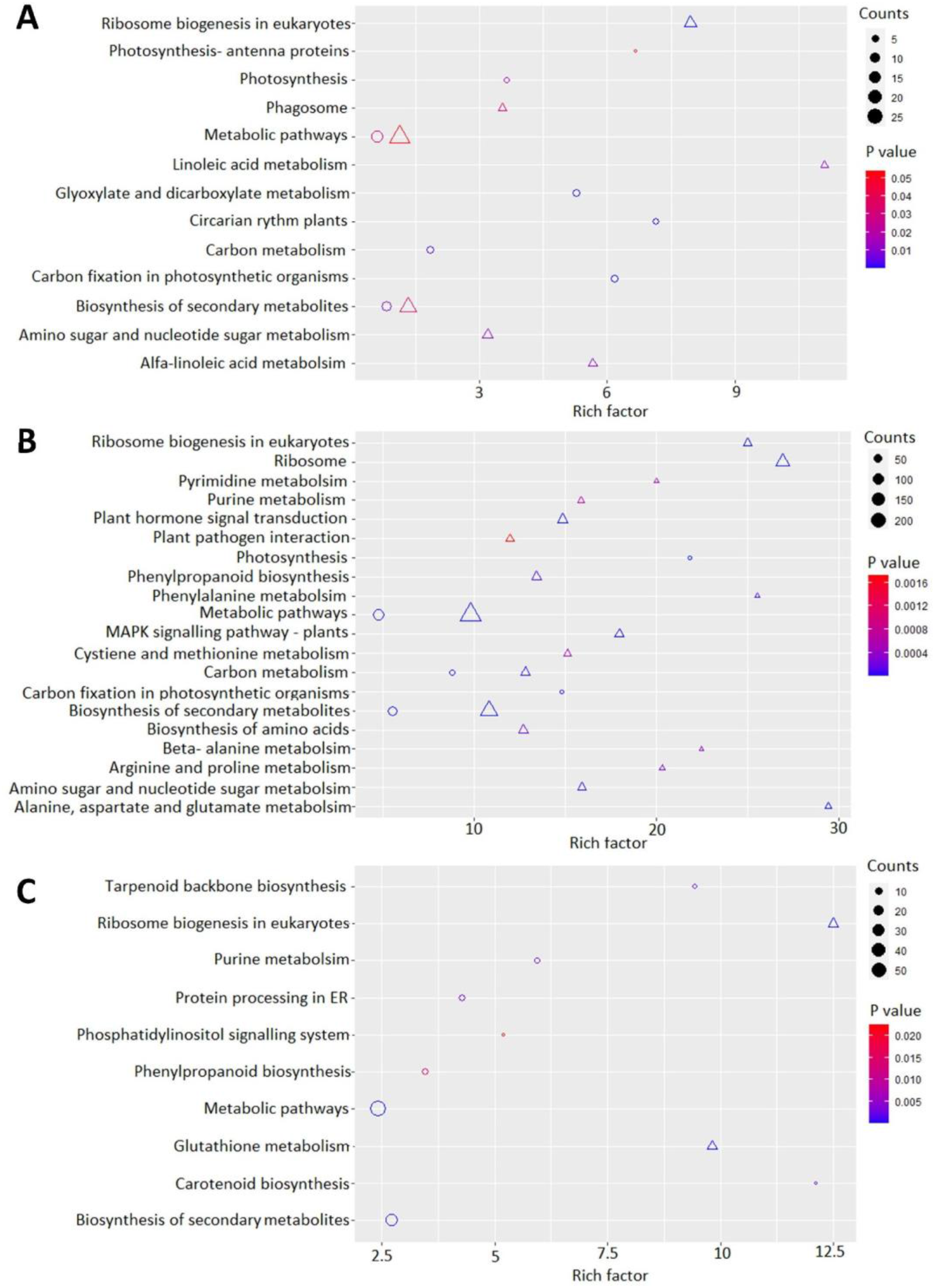
Pathway enrichment analysis. Dot plot showing top 20 significant regulatory pathways in rice response to *R. solani* infection at three Time Points i.e., day 1 (A), day 2 (B) and day 5 (C). Here, the size, shape and color represent the number of genes belonging to a particular pathway, pathway expression type (triangle represents up-regulation and circle represents down-regulation) and significance level respectively. Rich factor is the ratio of the number of genes in our DEG list annotated in a particular pathway to the total number of genes present in that pathway.

It was observed that 1^st^ and 2^nd^ day samples had more pathways in common as compared to 1^st^ and 5^th^ day. To gain deeper insights into gene level involvement, we checked the common genes within these overlapping (up-regulated and down-regulated) pathways appearing in day 1 and day 2 samples. Genes Os02g0218700/ AOS3, Os03g0179900/ LOX2 and Os08g0509100/ LOX8 belonging to ‘α-Linolenic acid metabolism’ and ‘Linoleic acid metabolism’ were found to be up-regulated and expressed commonly in day 1 and day 2 samples.

The pathway terms ‘Carbon fixation in photosynthetic organisms’, ‘Carbon metabolism’, ‘Glyoxylate and dicarboxylate metabolism’, ‘Photosynthesis’ and ‘Photosynthesis - antenna proteins’ were found to be down-regulated and expressed commonly in day 1 and day 2 samples suggesting compromised carbon fixation and photosynthesis in rice under *R. solani* infection conditions. Five of the genes belonging to the ‘Carbon fixation in photosynthetic organisms’, ‘Carbon metabolism’ and ‘Glyoxylate and dicarboxylate metabolism’ pathways i.e., Os12g0207600, Os01g0791033, Os05g0427800, Os10g0356000 and Os05g0496200 were found to be differentially expressed at both the Time Points i.e., day 1 and day 2. Among the five genes, Os05g0496200 codes for 3-phosphoglycerate kinase and the rest for the RuBisCO large subunit.

Pathways such as ‘MAPK signaling pathway’, ‘Phenylpropanoid biosynthesis’ and ‘Phenylalanine metabolism’ are also significant as their roles in rice in response to *R. solani* have been previously well documented (Karmakar et al., 2019; Ghosh et al., 2017; Zhang et al., 2017). Our results are in line with previous studies that also suggested their alteration in rice after *R. solani* infection. The pathways ‘Phenylalanine metabolism ‘and ‘Phenylpropanoid biosynthesis mostly consist of Peroxidases and PAL genes.

### Transcription Factors and families induced in rice in response to *R. solani* infection

Transcription Factors play crucial roles as either activators or repressors of genes in specific ways and are involved in a wide range of biological processes including plant defense against stressors (Smaczniak et al., 2012). TFs of rice spanned across 63 families of which TFs belonging to 39 families were found to be expressed in the present study. A total of 24 (13 up-regulated and 11 down-regulated), 212 (149 up-regulated and 63 down-regulated) and 76 (33 up-regulated and 43 down-regulated) TFs were found to be differentially expressed at 1^st^, 2^nd^ and 5^th^ dpi, respectively (Fig. 4A) (Supplemental Data Set 5). Of the 39 expressed TF families, 7 TF families showed significant change in expression after *R. solani* infection (Supplemental Table 2). It was observed that for each of the 7 TF families, the number of up-regulated TFs outnumbered the number of down-regulated TFs except for MYB-related family TFs expressed on day 5, where an opposite trend was observed.

**Figure 4.**
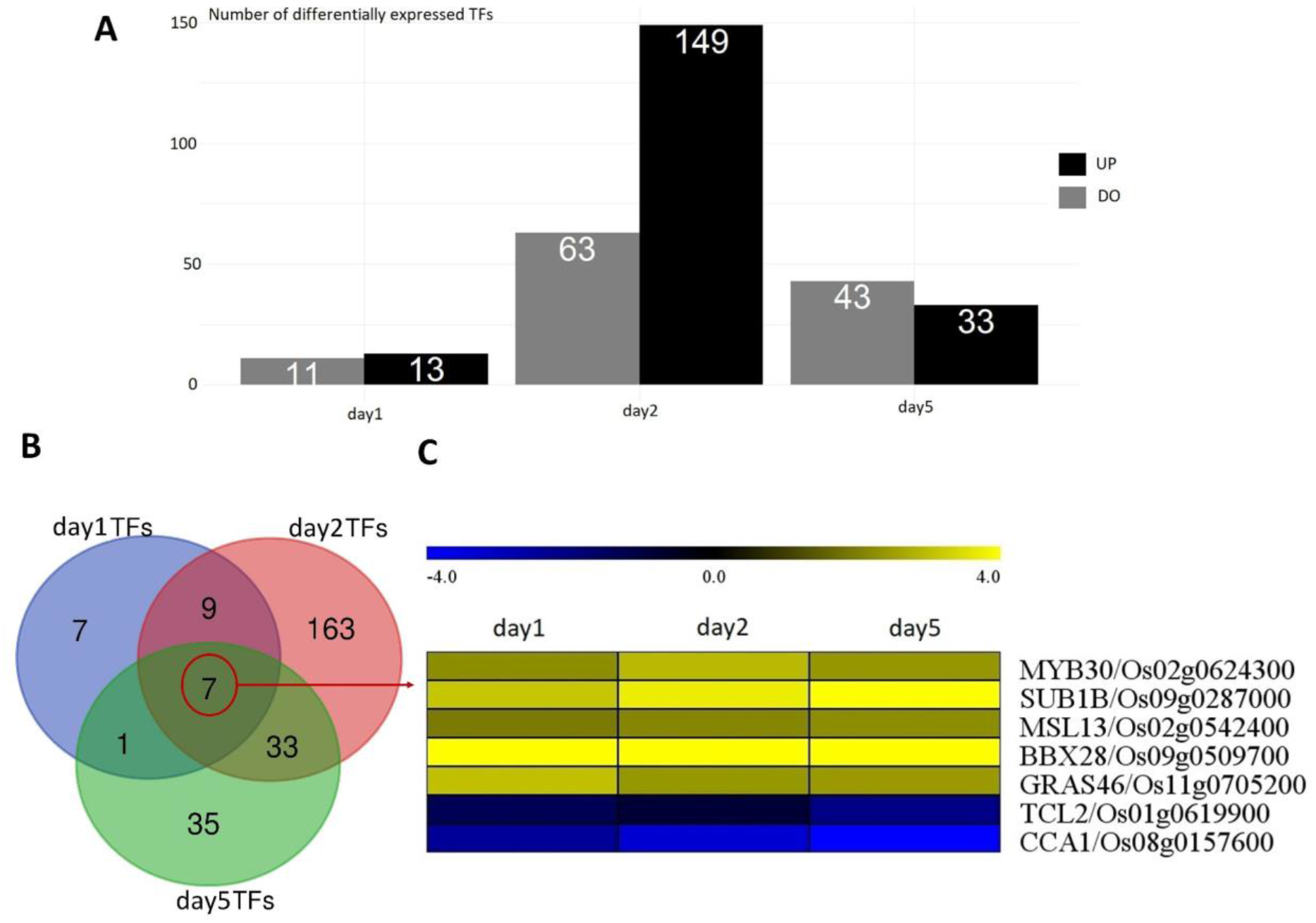
Rice Transcription Factors (TFs) responsive to *R. solani*. A, Bar graph showing the number of TFs differentially expressed in rice at 1, 2 and 5^th^ day Time Points after *R. solani* infection. B, Venn Diagram showing 7 TFs that were commonly expressed in all the three Time Points. C, Heatmap showing the expression profile of the 7 TFs that were found to be commonly expressed at all the three Time Points.

Further, we checked the TFs differentially expressed as common to all the three Time Points (Figure 4B). There were a total seven of them i.e., MYB30, SUB1B, MSL13, BBX28, GRAS46, TCL2 and CCA1 and all of them maintained uniform expression patterns with increasing Time Points suggesting their crucial role in rice responses to *R. solani* infection. Of the seven commonly expressed TFs, five were up-regulated and only CCA1, TCL2 were found to be down-regulated at all the Time Points (Fig. 4C).

### Topological rewiring of Transcriptional Regulatory Networks in rice during *R. solani* infection and identification of regulatory hubs

Biological network studies help in delineating how cells respond to a particular environmental condition like stress. In order to gain a systems level insight into the molecular mechanism in the rice plant post infection, we have constructed TF-TG regulatory networks for all the three Time Points using a machine learning approach (Fig. 5). Here, the Target Gene (TG) represents genes and TF represents Transcription Factors differentially expressed in each Time Point.

**Figure 5.**
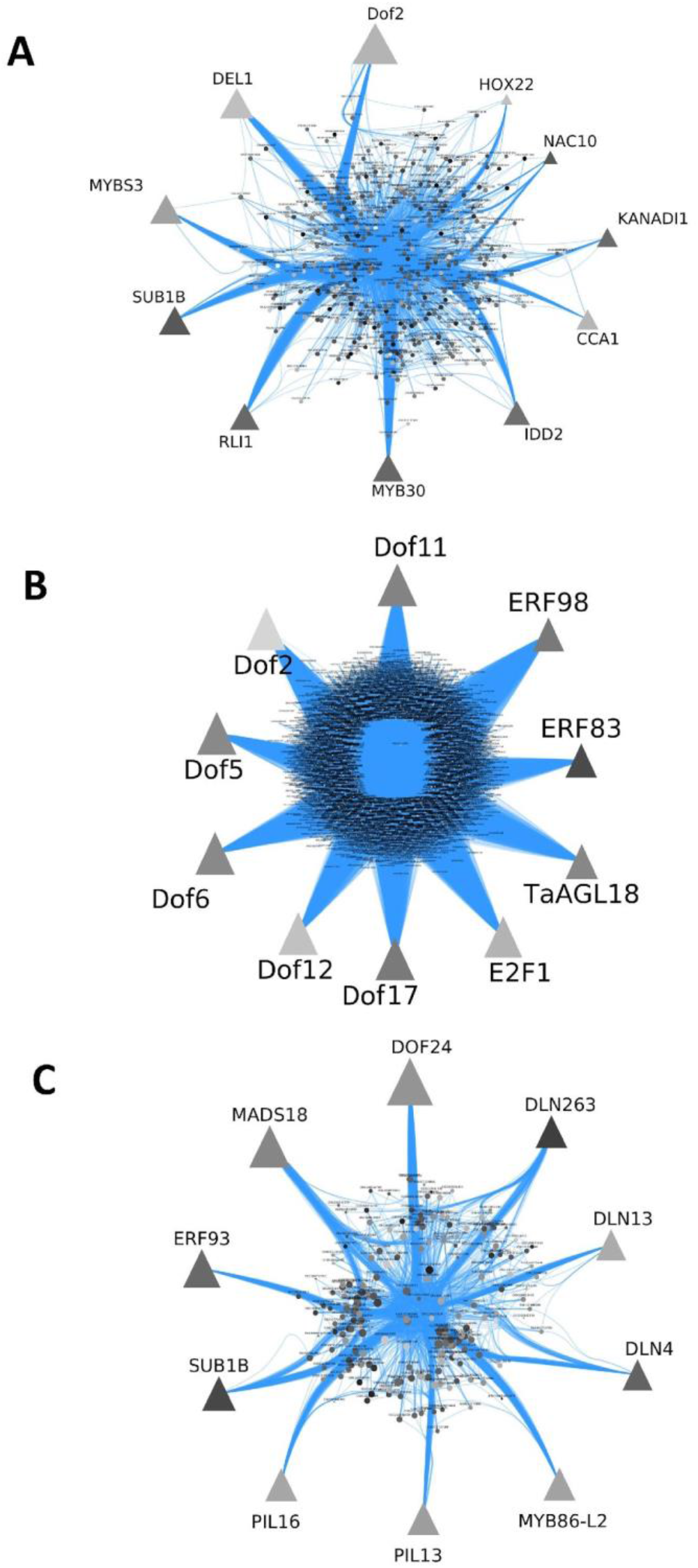
Topology of the Transcriptional Regulatory Networks (TRNs) in rice during *R. solani* infection. Transcription Factor – Target Gene (TF-TG) network of day 1 (A), day 2 (B) and day 5 (C) samples of rice after *R. solani* infection generated using motif scanning method. Here, the Target Gene (TG) represents genes and TF represents Transcription Factors differentially expressed in each of the Time Points.

There was a high variance among the sizes of the networks (Supplemental Table 3). Out of the three networks, the TF-TG network of day 2 was the largest in terms of number of edges and nodes. On analysing the topological parameters, it was found that the degree distribution of the networks did not follow a Poisson distribution. Hence, the constructed networks were not random in nature. A power-law was fit into each degree distribution and a goodness-of-fit test was calculated. The test was performed by a bootstrapping procedure of 1000 simulations. The result showed that only the degree distribution of the TF-TG network of day 2 could be explained by a power-law. In addition to being scale-free, the network is negatively assortative. This makes the network more robust than the other day 1 and day 5 networks.

Taking into consideration the parameter of ‘degree of centrality’, we identified 10 most highly connected TFs and marked them as regulatory hubs for each Time Point as shown in Supplemental Table 3. A dense TF regulatory network was obtained for day 2 with 2937 nodes and 163841 edges. Three of the TFs (marked with * in Supplemental Table 3) i.e. Os09g0287000/ SUB1B, Os02g0624300/ MYB30 and Os08g0157600/ CCA1 identified as regulatory hubs from the TF-TG network were also expressed as common to all the three samples (Fig. 4C). SUB1B and MYB30 showed consistency in maintaining higher expression levels whereas CCA1 was down-regulated throughout all the three Time Points.

### Putative Resistant Genes in Rice against *R. solani* and benchmarking

In order to benchmark and elucidate the role of putative *R. solani* resistant genes, our dataset was compared to a comprehensive and non-redundant list of 48 putative rice Resistant Genes (RGs) found in the three major diseases of rice i.e., Bacterial Blight (BB), Rice Blast (RB) and Sheath Blight (SB) disease. The non-redundant list was prepared by integrating data collected from PRGdb 3.0 database (Osuna-Cruz et al., 2018) (for BB and RB disease) and manual curation of literatures to find genes conferring resistance against SB owing to non-availability of any dedicated repository for SB RGs (Supplemental Data Set 6). Out of the genes, ten of the DEGs from this study matched with public data and were termed as Putative Resistant genes (Supplemental Figure S1). Out of them, 9 genes i.e., PR4, Os11g0569500/ XA21 (59.6% similar), Os11g0569300/XA21 (59.6% similar), 2H16, PGIP1, ACS2, OXO4, RCH10, PIT, chi11 were found to be up-regulated excluding 2H16. Additionally, majority of these Putative Resistant genes found their expression only on day 2 suggesting rice defense response to be stronger at this Time Point.

## DISCUSSION

### Transcriptional response of rice to *R. solani*

Transcriptome studies using RNA-Seq or microarray provide systems level insights into key genes, pathways and differentially regulated processes in a specific condition. The present study used a Time-Series RNA-Seq analysis to decipher the transcriptome responses of rice plants upon *R. solani* infection. We have identified 428, 3225 and 1225 DEGs in day 1, 2 and 5 samples, respectively. At every Time Point, the number of up-regulated genes always outnumbered the down-regulated genes. Day 2 sample showed the largest number of DEGs, which indicated that defense response of rice plants to *R. solani* was at the peak on 2nd dpi. Furthermore, identification of the genes belonging to the common biological processes that were expressed differentially at all the three Time Points (Supplemental Table 1) presents new avenues to identify SB tolerant genes in rice plants. Some of such processes have been described here under.

### JA biosynthesis and MAPK signaling pathways are key role players in rice plants during *R. solani* infection

Jasmonic acid signaling plays a central role in responses to necrotrophic pathogens by activating a plethora of pathogenesis-related (PR) genes (Tamaoki et al., 2013). Our study showed the induction of JA signaling in rice during *R. solani* infection. For example, the genes Os02g0218700/ AOS3, Os03g0179900/ LOX2 and Os08g0509100/ LOX8 belonging to ‘α-Linolenic acid metabolism’ and ‘Linoleic acid metabolism’ were found to be up-regulated and expressed on day 1 and day 2. Allene Oxide Synthase-AOS3 has already been identified as a candidate hub gene in Lemont rice variety responding to *R. solani* (Zhang et al., 2018). A recent study also showed that the expression levels of LOX-lipoxygenase were higher in SB resistant rice as compared to susceptible ones (Sayari et al., 2014). The genes AOS3, LOX2 and LOX8 are key participants in JA biogenesis and our results indicate that JA signaling pathway is activated in rice upon *R. solani* infection.

‘MAPK signaling pathway’ is also an important signaling pathway in rice plants in initiating defense responses to *R. solani* infection. We have identified 30 MAPK signaling pathway related genes that were up-regulated after *R. solani* infection (Fig. 3B and Supplemental Data Set 3). Mitogen-activated protein kinases (MAPK) are small molecules that take part in downstream signaling of receptors triggered by external stimulus like Pathogen/ Microbe-Associated Molecular Patterns (PAMPs/ MAMPs) and pathogenic effectors leading to the activation of numerous defense response mechanisms like biosynthesis of defense related hormones, ROS generation, stomatal closure, induction of PR genes and Hypersensitive Response (HR) leading to cell death (Meng and Zhang., 2013). About 75 putative MAPK are activated in rice upon different stress conditions (Rao et al., 2010; Kumar et al., 2008). A recent study also showed that MAPK6 was highly expressed in transgenic rice plant during SB disease condition (Karmakar et al., 2019).

### Alteration of CYPs, Peroxidases and PAL after *R. solani* infection

Cytochrome P450s (CYPs) are oxidoreductase enzymes that play important roles in biosynthesis of secondary metabolites, antioxidants, plant hormones etc. Additionally, they are also involved in plant defense responses (Pandian et al., 2020). An earlier study showed the role of CYP78, a plant specific CYP450 in imparting resistance to *R. solani* challenged rice plants by the overexpression of CYP78A encoding BSR2 (Broad Spectrum Resistance) gene (Maeda et al., 2019). In the present study, the gene ‘CYP51H4/ Os02g0323600’ a member of both the ‘Metabolic pathways’ and ‘Biosynthesis of secondary metabolites’ pathways was found to be strongly downregulated at all the Time Points post infection.

The secondary metabolic pathways ‘Phenylpropanoid biosynthesis’ and ‘Phenylalanine metabolism’ (Fig. 3B) were found to be up-regulated upon *R. solani* infection in early infection stage. Most of the genes in the pathways of ‘Phenylalanine metabolism ‘and ‘Phenylpropanoid biosynthesis include Peroxidases and PAL. Previous studies suggest that Peroxidase and PAL genes were activated upon *R. solani* infection in rice, with PAL acting as a constituent of multi-component coordinated defense response mechanism against the pathogen (Bera et al. 1999), (Mutuku et al., 2012). Phenylpropanoid biosynthesis is known to be involved in the formation of secondary metabolites like phytoalexin, lignin and phenolic substances. Both Phenylpropanoid and PAL genes were activated in TeQing a moderately resistant rice cultivator upon *R. solani* infection (Zhang et al., 2017). Induction of Peroxidases and PAL genes might constitute a protection strategy initiated by rice to reinforce its cell wall cross linking thereby preventing further invasion by fungal hyphae. These results suggest that Peroxidases and PAL metabolism have significant role in early infection stage in the present case.

### Photosynthesis machinery is down-regulated after *R. solani* infection

Plants tend to compromise their primary processes such as photosynthesis and reproduction upon exposure to stressed conditions in order to allocate more resources to activate defense, stress and secondary metabolic processes (Barah et al., 2015). For example, five of the genes belonging to the ‘Carbon fixation in photosynthetic organisms’, ‘Carbon metabolism’ and ‘Glyoxylate and dicarboxylate metabolism’ pathways i.e., Os12g0207600, Os01g0791033, Os05g0427800, Os10g0356000 and Os05g0496200 were found to be commonly down-regulated at day 1 and day 2 Time Points. Out of these five genes, Os05g0496200 codes for 3-phosphoglycerate kinase and the rest code for the RuBisCO large subunit. RuBisCO is the primary enzyme for carbon fixation in plants. Previous studies have shown that rice, *Arabidopsis*, tomato and soyabean showed decreased RuBisCO amount and function upon exposure to drought stress (Majumdar et al., 1991). For the pathway ‘Photosynthesis’, the gene Os01g0938100 encoding Psb28 was found to be common to both day 1 and day 2 samples. Psb28 is a constituent of photosystem II and it has been shown that the disruption of psb28 gene in rice encourages pale green type morphology in rice (Jung et al., 2008). Recently, GC-MS analysis of *R. solani* susceptible rice cultivar, Pusa Basmati-1 (PB1) found deformed chloroplast ultrastructure and reduced photosynthetic efficiency after infection (Ghosh et al., 2017). Our study results also strongly suggested that Photosynthesis machinery was compromised upon *R. solani* infection.

### WRKY, NAC, ERF and MYB family Transcription Factors are induced after *R. solani* infection

Our analysis identified that the expression patterns of WRKY, NAC, ERF and MYB TFs were significantly altered in rice plants in response to the treatment with *R. solani*. WRKY TFs are considered as central elements for rice resistance to various pathogens (Khong et al., 2008). In the recent past, WRKY TF genes have also been shown to be the key role players in rice defenses against *R. solani*. To our knowledge, so far WRKY4, WRKY13, WRKY80 and WRKY30 have been identified to impart resistance in rice to *R. solani* infection (Wang et al., 2015a; Lilly et al., 2019; Peng et al., 2016; Peng et al., 2012). Out of the 27 WRKY TFs identified to be expressed in our study, 26 were up-regulated at day 2 and day 5 Time Points (Supplemental Table 2). A recent study also suggested that the expression of WRKY30 gene enhances endogenous JA accumulation in transgenic rice thereby imparting resistance to SB disease (Peng et al., 2012). As the JA biosynthetic genes i.e., AOS3, LOX2 and LOX8 were also found to be up-regulated in our analysis, it can be speculated that the overexpression of WRKY TFs might have led to enhanced JA synthesis in rice after *R. solani* infection. Various TFs belonging to NAC, MYB and ERF TF families were more up-regulated after *R. solani* infection (Supplemental Table 2). In line with these observations, another recent study also suggested the involvement of MYB and NAC TFs in rice SB responses (Peng Yuan et al., 2020). To summarize, WRKY, NAC, ERF and MYB TFs are highly induced in rice upon *R. solani* infection.

### Transcriptional Regulatory Network identifies SUB1B, MYB30 and CCA1 as key response signatures

Most TFs bind to recognition sites called *cis*-regulatory elements in the promoter regions of target genes (TGs) leading to subsequent activation or deactivation of the corresponding gene. Differential re-wiring of the TF-TG associations provide a genome-scale overview of the transcriptional re-programming (Lim and Xie, 2018). The topology of the transcriptional regulatory networks can be used for identifying hub genes. The key hub Transcription Factors identified from our analysis were SUB1B, MYB30 and CCA1. These three hub TFs constitute rice core responses to *R. solani* infection. All of them showed uniform expression during all three Time Points. SUB1A is an ERF family TF responsible for both drought and submergence tolerance in rice by repressing the Gibberellic Acid response pathway through the up-regulation of DREB1s and AP59. It also regulates ROS production to prevent chlorophyll and oxidative damage during re-oxygenation (Fukao et al. 2010). SUB1B is an allele of SUB1A and might have a similar role in regulating ROS production in the cell. SUB1B might also be involved in defense against *R. solani* apart from flood tolerance.

MYB30 TF is a microbe-associated molecular pattern (MAMP) responsive TF, which enhances resistance in rice against bacterial and fungal pathogens by producing more hydroxycinnamic acids (HCAAs) (Kishi-Kaboshi et al., 2018). HCAAs are phenylpropanoids made from phenylalanine. They bind to components of the cell wall and facilitate cell wall cross-linking (de Oliveira et al., 2015). From our pathway enrichment analysis, it is evident that ‘Phenylalanine metabolism’ and ‘Phenylpropanoid biosynthesis’ pathways are crucial parts of rice responses to *R. solani*. Therefore, MYB TF family members like MYB30 may be involved in the production of more HCAAs. MYB30 is also a negative regulator of cold tolerance in rice by suppressing β-amylase activity and hence, reducing Maltose production that is responsible for cell membrane protection against cold stress (Lv et al., 2017).

Circadian Clock Associated 1 (CCA1) is a component of Morning Element Loop of circadian cycle and reaches its peak during dawn. Circadian Clock Associated 1 (CCA1) and Timing of Cab Expression 1 (TOC1) are known as central circadian clock oscillators. Circadian clock system has been shown to regulate ROS homeostasis, Abscisic Acid stress response signaling and various R genes, and their targets in plants in response to various stressors. The cycle initiates in the presence of light the activation of CCA1 and gradually ceases by the evening due to higher TOC1 gene expression. As the cycle progresses from morning to evening, CCA1 decreases and TOC1 increases with its peak at dusk (Bhattacharya et al., 2017). In the present study, CCA1 identified as a hub TF was down-regulated and in contrast, its partner TOC1 was highly up-regulated at all the three-day Time Points suggesting the strong influence of *R. solani* on circadian cycle of the rice plant. Recently, it was shown that TOC1 and Abscisic Acid work together in a feedback loop and is important for plant response to drought stress condition (Legnaioli et al., 2009).

### SB resistance genes in rice

A list of 10 putative resistant genes were found to be expressed in our data, and seven of them belonged to SB resistance gene list. The gene Os11g0592200/ PR-4b belongs to the class II of the PR-4 family and is located in the vacuole. It has also been shown that rice PR-4b exhibits antifungal activity against *R. solani*, but the resistance mechanism underlying the antifungal activity is yet to be revealed (Zhu et al., 2006). Os11g0569500 and Os11g0569300 are *Xa21* homologues and share 59.6% similarity with *Xa21. Xa21* has always been a popular target for researchers studying rice resistance to Bacterial Blight. *Xa21* imparts broad spectrum resistance against Bacterial Blight disease by altering various energy distribution schemas and targeting phytohormones (Peng et al., 2015). The present study also suggests that *Xa21*, already known for BB resistance might also be important for rice resistance against SB disease. Recently, transgenic rice lines overexpressing 2H16 exhibited enhanced tolerance to SB disease (Li et al., 2013). In our study, Os06g0316000/ 2H16 was the only down-regulated SB resistant gene. The gene Os05g0104200/ PGIP1 has been reported to counter-attack polygalacturonases (PG) secreted by the fungal pathogens to degrade plant cell wall and rice overexpressing PGIP1 showed significant resistance to *R. solani* (Wang et al., 2015).

Ethylene biogenesis signaling pathway has been identified to confer resistance to various diseases along with the modulation of JA and SA pathways. Os04g0578000/ ACS2 1-aminocyclopropane-1-carboxylic acid synthase is a key gene in Ethylene biogenesis and its overexpression in transgenic rice lines imparted resistance to SB disease (Helliwell et al., 2013). The transgenic lines developed with Os03g0694000/ OXO4 and Os06g0726100/ chi11 combined cassettes exhibited significant resistance to SB disease. OXO4 imparts resistance by activating SAR by nullifying OA (Oxalic Acid) produced by *R. solani* leading to generation of hydrogen peroxide (Karmakar et al., 2016). Another transgenic rice line study using combined RCH10 and AGLU1 showed that their proteins were localized in chloroplast and combined expression of these two genes enhanced resistance to SB disease in rice (Mao et al., 2014). Os01g0149500 is a Pit gene homologue with 99.6% similarity with Pit. Pit gene is one of the resistance genes cloned and identified among 102 RGs against rice blast disease (Xiao et al., 2017). Our dataset contains 7 genes viz. PR4, 2H16, PGIP1, ACS2, OXO4, RCH10, chi11, which were reported to confer resistance to rice in SB disease condition.

## CONCLUSION

Using systems biology approaches, the study identified key molecular signatures that are altered in rice plant after *R. solani* infection at three Time Points. Our results have shown that rice plant differentially modulates its transcriptome landscape in response to *R. solani* attack and the response strategies during early and late infection phases share both common and unique signatures. Induction of defense related signaling pathways and several key molecules suggest activation of strong immune responses in rice upon *R. solani* infection and such redistribution of energies led to the down-regulation of photosynthesis machinery. *R. solani* infection caused differential re-wiring of transcriptional regulatory connections in rice and the connectivity patterns drastically varied as the infection progressed. The study also detected the differential expression of seven *R. solani* resistant genes. Further research is required to identify whether the genes identified to be strongly influenced by *R. solani* infection also impart tolerance against SB disease in rice. The knowledge gained from this study will help develop a better understanding of the disease and serve as a valuable resource for development of future SB tolerant rice varieties.

### Accession Numbers

Datasets generated from this study has been deposited in NCBI-SRA database with BioProject accession number PRJNA725619.

## Figure legends

**Figure 1**. Differentially expressed genes (DEGs) in rice after *R*. *solani* infection. A, Bar plot showing the number of DEGs (up-regulated and down-regulated) identified in rice at 1^st^, 2^nd^ and 5^th^ day Time Points after *R*. *solani* infection. B, Venn diagram showing up-regulated DEGs expressed as common in day 1, 2 and 5 samples. A total of 133 genes were found to be up-regulated as common in all the three Time Points. C, Venn diagram showing down-regulated DEGs expressed as common in day 1, 2 and 5. A total of 20 genes were found to be down-regulated as common in all the three Time Points.

## Supplemental Data

The following supplemental information is available.

**Supplemental Figure S1**. Expression profile of resistant genes in rice against Sheath Blight (SB), Rice Blast (RB) and Bacterial Blight (BB) diseases in our dataset.

**Supplemental Table 1**. Common genes of Biological Processes that were expressed at 1^st^, 2^nd^ and 5^th^ day Time Point of *R. solani* infection.

**Supplemental Table 2**. Altered expression of rice TF families at 1^st^, 2^nd^ and 5^th^ day Time Point of *R. solani* infection.

**Supplemental Table 3**. Hub Transcription Factors identified from Transcriptional Regulatory Networks (TRNs) at 1^st^, 2^nd^ and 5^th^ day Time Point of *R. solani* infection.

**Supplemental Data Set 1**. List of differentially expressed genes (DEGs) in rice plant at 1^st^, 2^nd^ and 5^th^ day of *R. solani* infection.

**Supplemental Data Set 2**. Altered Biological Process (BP), Cellular Component (CC) and Molecular Function (MF) terms in rice plant at 1^st^, 2^nd^ and 5^th^ day of *R. solani* infection.

**Supplemental Data Set 3**. Altered KEGG pathways in rice plant at 1^st^, 2^nd^ and 5^th^ day of *R. solani* infection.

**Supplemental Data Set 4**. Comparison of differentially regulated pathways in rice plant at 1^st^, 2^nd^ and 5^th^ day of *R. solani* infection.

**Supplemental Data Set 5**. List of Transcription Factors that were differentially regulated in rice plant at 1^st^, 2^nd^ and 5^th^ day of *R. solani* infection.

**Supplemental Data Set 6**. Resistant genes in rice against three major diseases i.e., Bacterial Blight, Rice blast, Sheath Blight from PRGdb 3.0 database and manual curation of literatures.

## ACKNOWLEDGMENTS

This research was supported by funding received from the Department of Biotechnology (DBT) (grant id - BT/PR24757/NER/95/843/2017) for the DBT-Twinning project-‘Integrative system biology approach to identify the molecular response signatures in Rice during concurrent biotic (*Rhizoctonia solani*) and abiotic (heat) stresses’ to Dr. Pankaj Barah. P.B.K acknowledges the award of NASI-Platinum Jubilee Senior Scientist by the National Academy of Sciences, India.

